# *WISP2/CCN5* gene knockdown in vitro and in vivo exhibits proliferation promotion of breast cancer through targeting Skp2 and p27Kip1

**DOI:** 10.1101/2020.01.29.924688

**Authors:** Yan Lv, Chang Zhang, Xiao Jiang Li, Shan Gao, Xu Zheng, Yan Yan Han, Chong Li, Qiang Geng

## Abstract

**Background:** Emerging evidence has demonstrated that WISP2/CCN5 is critically involved in tumorigenesis. However, the function of WISP2/CCN5 in breast cancer carcinogenesis is largely unclear.

**Methods:** we aim to explore the effects and potential mechanisms of WISP2/CCN5 on proliferation of breast cancer cells and carcinogenesis of breast cancer xenograft. Lentivirus vector with WISP2/CCN5shRNA was transfected into MCF-7, and breast cancer cells and xenograft were conducted. Effect of WISP2/CCN5 on growth and carcinogenesis of breast cancer cells and xenografts was evaluated by MTT assay and tumor volume. The relationship between WISP2/CCN5, Skp2 and p27Kip1 was detected in vitro and in vivo by RT-PCR at mRNA level and Western blotting at protein level.

**Results:** The result of MTT assay indicated that MCF-7 cell growth viability in WISP2/CCN5 gene knockdown group was significantly higher than negative vector group(*P*<0.05) or control group (*P*<0.05). It suggested that knockdown of *WISP2/CCN5* gene by shRNA lentivirus plasmid promoted proliferation of MCF-7 cells. The growth curves of breast cancer xenograft showed that xenografts in WISP2/CCN5 knockdown group grew more quickly than negative vector group(*P*< 0.05) or control group (*P*< 0.05). Subsequently, the results of RT-PCR and Western blotting revealed that *WISP2/CCN5* gene knockdown led to increased Skp2 and decreased p27Kip1 at mRNA and protein levels. WISP2/CCN5 exerts its inhibition on proliferation of MCF-7 cell line and suppressive functions on growth of breast carcinoma via regulation of Skp2 and p27Kip1at mRNA and protein levels. However, *WISP2/CCN5* gene knockdown resulted in loss of inhibition effect on MCF-7 and breast cancer.

**Conclusions:** Our findings suggest that WISP2/CCN5 could be a useful therapeutic strategy for the treatment of breast cancer through targeting Skp2 and p27Kip1.

## Introduction

Breast cancer is one of leading causes of cancer-related death in women worldwide. To explore the mechanism of occurrence and progression of breast cancer, to accurately treat breast cancer is of great significance. Recently, important roles of CCN5/WISP2 proteins have been recognized in breast cancer progression ^[1]^. WISP-2 (Wnt-1 induced secreted proteins −2, Wnt secretory protein 2) plays an important role in inducing cell apoptosis, inhibiting cell adhesion and metastasis ^[2]^. WISP2/CCN5 was reported to be upregulated in the mammary epithelial cells transformed by the Wnt-1 oncogene ^[3]^. WISP2/CCN5 was found to be directly regulated by the estrogen receptor in human breast cancer cells ^[4]^. One study further identified that WISP2/CCN5 expression was enhanced by serum and correlated with serum-induced cell proliferation in breast cancer cells, demonstrating that WISP2/CCN5 could enhance cell proliferation in breast cancer ^[5]^.

The F-box protein Skp2, a component of the Skp1-Cullin 1-F-box (SCF) E3 ubiquitin-ligase complex, has been shown to regulate cellular proliferation, carcinoma progression and metastasis by targeting several cell cycle regulators for ubiquitinationr^[6]^. P27^KIP1^ is a negative regulator of cell cycle that plays an important role in tumor suppression. Reduction of p27^KIP1^ secondary to enhanced Skp2-mediated degradation is involved in atypical hyperplasia and tumorigenesis. ^[7]^.

In present study, we intend to find out the relationship between *WISP2/CCN5* gene knockdown and proliferation and progression of breast cancer in vitro and in vivo, and those underlying molecular mechanisms.

## Materials and Methods

### Cell line and cell culture

Human breast cancer cell line MCF-7 was purchased from KGI biotechnology co., LTD, Jiang Su, China and cultured in Dulbecco’s modified Eagle’s medium (DMEM, Gibco, USA) containing 10% fetal bovine serum (FBS, Gibco, USA) and 1% penicillin-streptomycin solution (Gibco, USA) in a incubator at 37°C and a humidified 5% CO_2_ atmosphere. MCF-7 passage cell line were prepared using 0.05% trypsin solution (Invitrogen, Carlsbad) and seeded in 96 well tissue culture plates. Expression plasmis and establishment of WISP2/CCN5 stably knockdown MCF-7 breast cancer cell line.

WISP2/CCN5 shRNA lentivirus plasmid was obtained from Hesheng Gene Biotechnology Co.Ltd, Beijing, China. One of WISP2/CCN5 shRNA sequence were designed according to targeted human WISP2/CCN5 gene and anchored with lentivirus plasmid in order to interfere the transcription of WISP2/CCN5 gene. The shRNA sequences were as follows: 5’GCCTACACACACAGCCTATATCGAAATATAGGCTGTGTGTGT FTAGTCC3’;

Another shRNA sequence was designed and combined with lentivirus plasmid so as to be used as null vector, sequences as follows: 5’CCTAAGGTTAAGTCGCCCTCGCCGAAGCGAGGGCGACTTAA CCTTAGG 3’

MCF-7 cell line was incubated in transfection solution of WISP2/CCN5 shRNA supplemented with 5μg/ml polybrene (transfection fortifier) in order to establish WISP2/CCN5 stably knockdown MCF-7 breast cancer line. Eight hours later, MCF-7cell line was treated with DMEM culture medium in the place of transfection solution.

### Cell viability assay

MTT assay was performed to assess cell proliferation. The MTT Kit was purchased from Beijing Solarbio Science and Technology Co., Ltd, and executed as kit specification. When cell density was about 70%, digested and adjusted the cell density at 1×10^4^ cells/ml, inoculated 180μl/well into 96-well plate, set up 6 replicate wells, including *WISP2/CCN5* gene knockdown cell line, null vector cell line and control cell line. After 24 hours, added 90μl of DMEM and 10 μl of 3-[4, 5-dimethylthiazol-2-yl]-2, 5 diphenyltetrazolium bromide (MTT) solution per well, continued to incubated at 37°C for 4 hours. Carefully aspirated the supernatant, added110 μl Formazan per well, shaked 96-well plate for 10 minutes in ahorizontal shaker in order to dissolve the dye, and measured OD value at 490 nm.

### Experimental animals

The experimental animals were purchased from SPF (Beijing) Biotechnology Co., Ltd., Beijing, China, and housed animal care facilities in Institute of Radiation Medicine Chinese Academy of Medical Sciences, Tianjin, China. *WISP2/CCN5* gene knockdown conditional transgenic mice were generated at Institute of Radiation Medicine Chinese Academy of Medical Sciences. The female nude mice, 6 weeks of age, 18-20g of weight, were chosen as experimental animals. The mice were randomized into three groups, 15 nude mice each group and were treated differently. The Institutional Animal Care and Use Committee (IACUC) were from Ethics Committee of Tianjin University of TCM, China.

### Establishment of genetically engineered mice model

The stably WISP2/CCN5shRNA knockdown MCF-7 cell line, negative vector MCF-7 cell line and control MCF-7 cell line were cultured and ogarithmic growth phase cells were digested and resuspended in PBS, adjusted cell density to 2× 10^6^ cells/ml.

The nude mice were injected 0.2ml of 2×10^6^ cells/ml WISP2/CCN5shRNA knockdown MCF-7 cell line, negative vector MCF-7 cell line and control MCF-7 cell line in right nipple region subcutaneously. The injection was executed once a day. After about 2 weeks, mice were executed, and tumors were taken, the tumor volume were determined.

### Xenografts volume measurement

Tumor size was measured daily using a caliper, and the tumor volume was determined with the standard formula: L× W^2^×0.5^2^, where L is the longest diameter and W is the shortest diameter. The growth curve of xenograft delineated the tendency of WISP2/CCN5 gene knockdown xenograft transformation during 14 days.

### Quantitative RT-PCR analysis

In order to find out the mechanism of *WISP2/CCN5* regulating MCF-7 proliferation in mRNA level, quantitative RT-PCR was performed to explore mRNA level of *WISP2/CCN5*, *Skp2* and *p27Kip1*in both MCF-7cells and xenografts.

RNA in vitro MCF-7 cell line and in vitro breast carcinoma tumor tissue were isolated using RNA prep pure cell/Bacteria kit and RNeasy reagents (TIANGEN BioTech Co.LTD, Beijing, China). Concentration and purity of RNA were detected by nucleic acid analyzing machine (Thermo Fisher, Germany). Reverse-transcription was performed using FastKing RT Kit (TIANGEN BioTech Co.LTD, Beijing, China) according to the manufacturer’s instructions. First, gDNA was degraded in a reaction system, including 5× gDNA Buffer 2 μ l, total RNA, RNase –Free ddH_2_O up to 10 μ l at 42°C for 3 min. Then, Reverse transcription was carried out using FQ -RT Primer Mix 2μl, FastKing RT Enzyme Mix 1 μ l, 10× King RT Buffer 2μ l, Rnase-Free ddH_2_O up to 10μ l, and mixed with gDNA reaction system for 40 cycles of 30 sec at 60°C. Amplifications of WISP2/CCN5, Skp2 and p27KIP1 were then performed using the cDNA template, SYBR Premix Ex TaqTM, 10μmol forward and reverse primers, and dH_2_O in a 25 μl volume using a QuantiTect SYBR Green PCR kit(TIANGEN BioTech Co.LTD, Beijing, China).Transcripts were quantified with β-actin as an internal standard. The sequences for the primers were as follows:

CCN5 forward, 5’CTGGGCTGATGGAAGATGGT3’, CCN5 reverse,3’TGTGTGTGTAGGCAGGGAGTG 5’; Skp2 forward, 5’ATGGACCAACCATTGGCTGAA3’, Skp2reverse,3’ACACTGAGACAGTATGCCGTGGAG5’; p27Kip1 forward, 5’CAAATGCCGGTTCTGTGGAG3’, p27Kip1 reverse, TCCATTCCATGAAGTCAGCGATA5’.

The cycling parameters were as follows: 30 s at 95°C,5 s at 95°C, and then 40 cycles of 30 s at 63.3°C (forp27KIP1), 64.6°C (for Skp2) or 60°C (for WISP2/CCN5) and 30s at 60°C.

### Western blot analysis and antibodies

Western blotting was performed according to the established methods. The breast tumor tissues were cut into small pieces. Cells were rinsed twice with ice-cold phosphate buffered saline (PBS, pH7.4), lysed in a cooled buffer. Subsequently, PMSF was added into xenograft tissues homogenate or cells, and pulverized. Then the lysates were boiled in a water bath for 10 minutes, and centrifuged at 12,000 g for 10 minutes at 4 °C.

The protein concentrations in the supernatant were measured by BCA kit (Solarbio, Beijing,China). Equal amounts of proteins were separated by 12% SDS-polyacrylamide gel electrophoresis (SDS-PAGE) and transferred to nitrocellulose membranes and immunoblotted with primary antibody overnight. After washing, the membrane was incubated with HRP-conjugated anti-mouse or anti-rabbit secondary antibody (Amersham, Germany). As appropriate, and the target proteins were visualized by ECL Plus Western Blotting Detection System.

The primary antibodies are anti-WISP2/CCN5 (1:1000, Arigo, USA), anti-Skp2 (1:500, Affinity, China) and anti-p27Kip1 (1:150, Affinity, China). For quantitative analysis, the films were scanned and analysis was performed using Image J software.

### Statistical analysis

The statistical analysis was performed using the Graph Pad Prism 4 (Graph Pad Software, Inc, La Jolla, CA, USA) and PASS^15.0^ software. Results are shown as mean ± S.D. Means between the groups were calculated and compared among or within variants using a two-sided Student’s t-test. Pearson Correlation was used to analyze the correlation between WISP2/CCN5 and Skp2 and p27Kip1. P value of <0.05 was considered statistically significant.

## Results

### Knockdown of *WISP2/CCN5* gene promoted the proliferation of MCF-7cell line

To validate the effect of WISP2/CCN5 gene knockdown on MCF-7 cells proliferation, transfection of shRNA targeted WISP2/CCN5 gene was executed. Figure 1 showed that MCF-7 was transfected with WISP2/CCN5 shRNA(A) and negative vector(B). There was no significant difference between two groups in fluorescence intensity and transfected cell percentages. MCF-7 was transfected by recombinant WISP2/CCN5shRNA lentivirus plasmid (figure 1A) and negative lentivirus plasmid (figure 1B) with GFP fluorescent clusters.

**Figure 1A.**
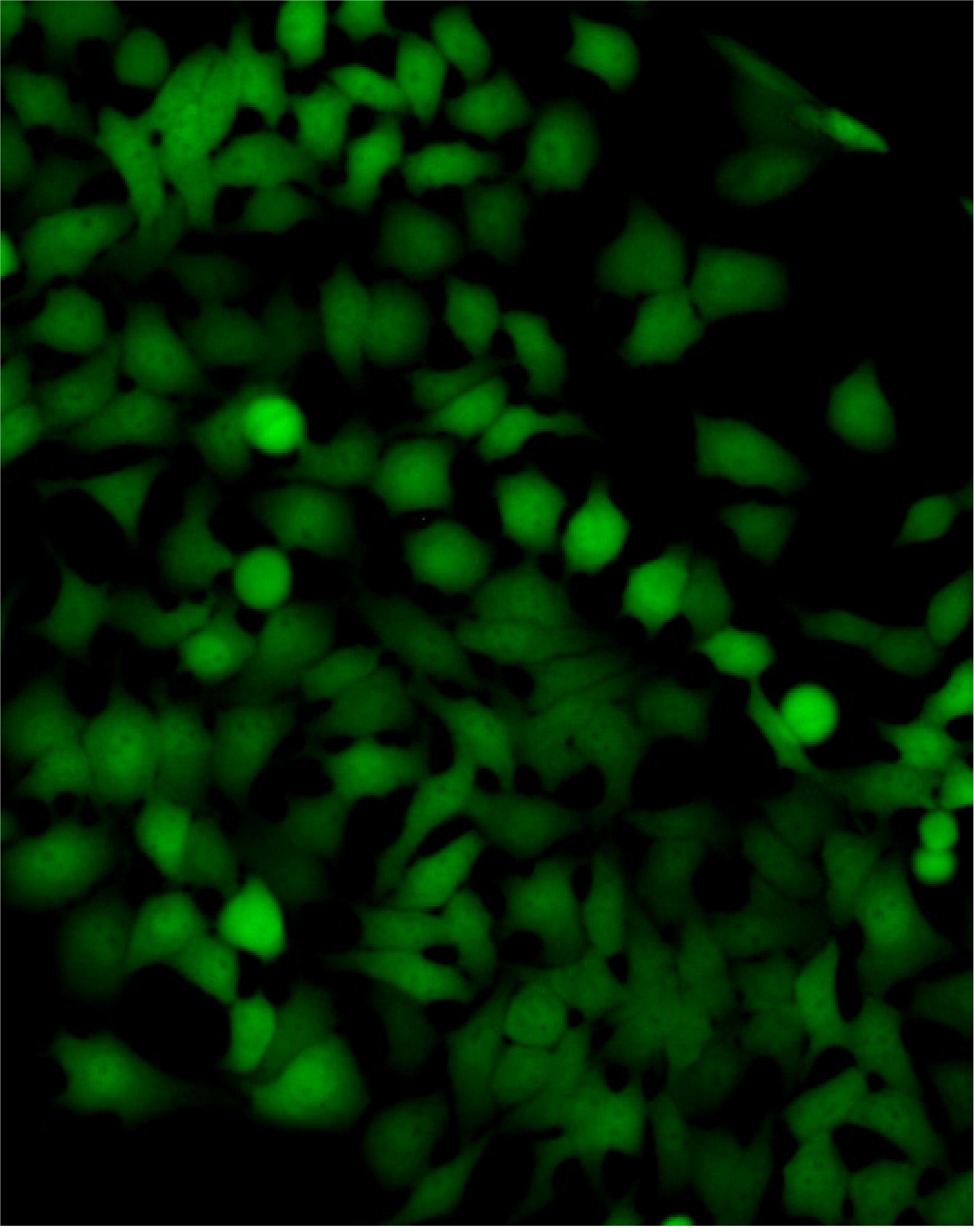

**Figure 1B.**
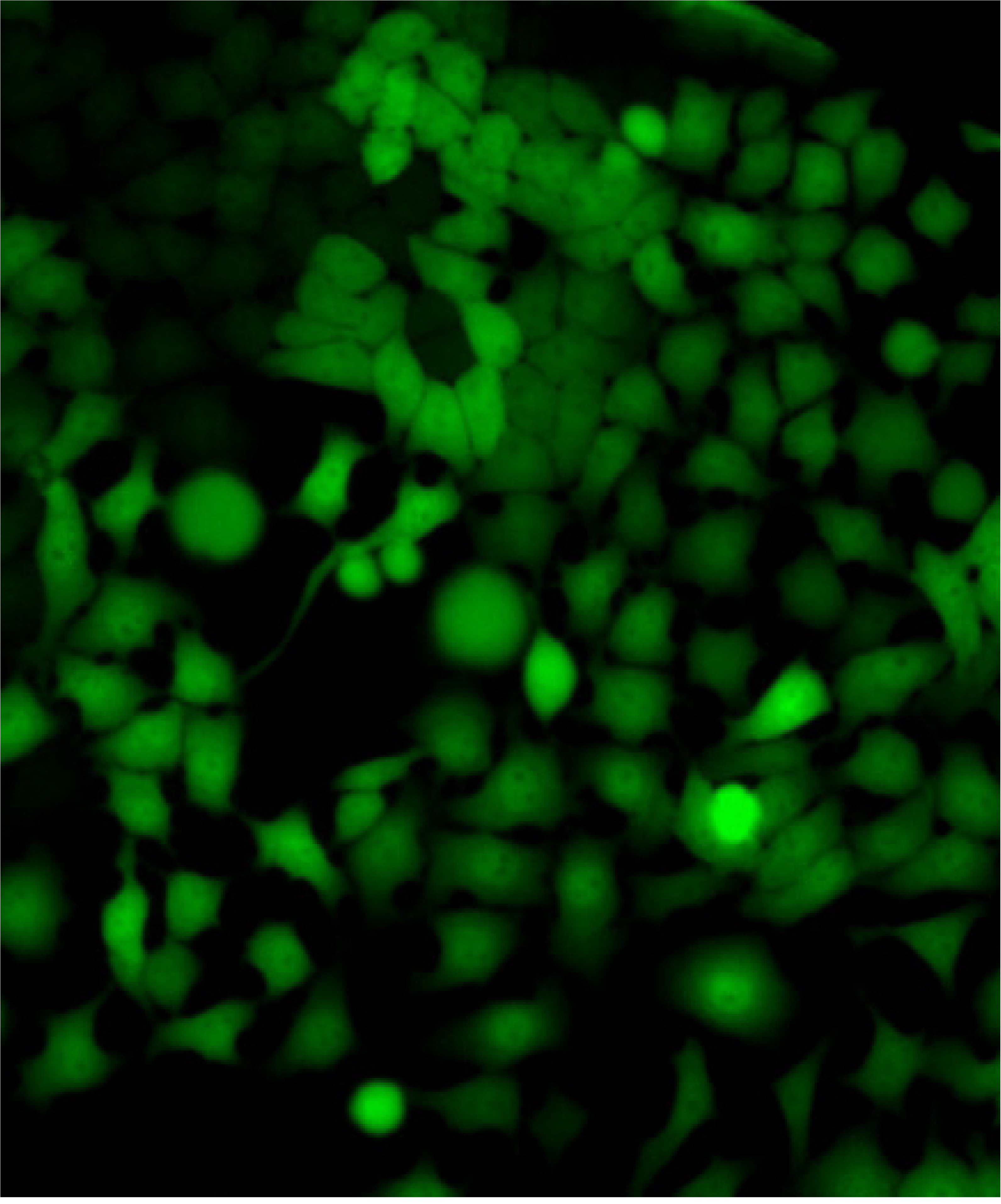

Tansfected cell relative viability was determined by MTT assay, and it demonstrated that shRNA transfection targeted WISP2/CCN5 gene leaded to remarkable proliferation of MCF-7 cell line, which exhibited higher proliferation and colony formation compared with control and negative vector cells (Figure 2). In addition, cell relative viability = gene knockdown group OD value/control group OD value.

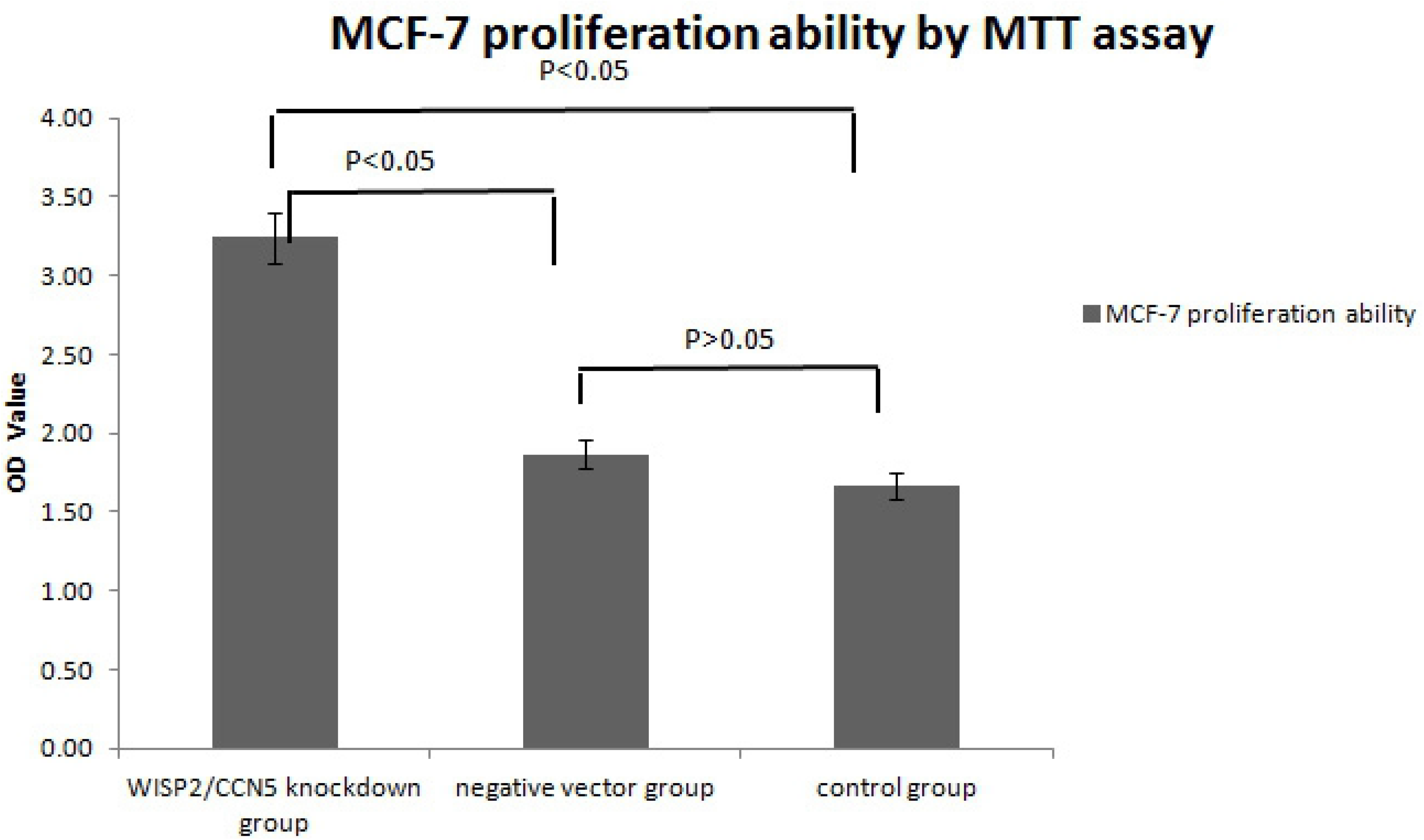
MTT analysis.

These results revealed that the viability of MCF-7 cells was decreased with dose of WISP2/CCN5shRNA lentivirus vectors compared with control group (*P*<0.05) or negative vector group (*P*<0.05).

The promotion on MCF-7 viability with dose of null lentivirus vectors was not markedly compared to control group (*P*> 0.05). The results were showed in Figure 2.

### Knockdown of *WISP2/CCN5* gene accelerated the growth of breast cancer xenograft

In order to explore the role of WISP2/CCN5 gene in genesis of breast carcinoma, the volume of engineered mouse xenograft was measured and compared between WISP2/CCN5 gene knockdown group and null vector group or control group.

The results of animal experiment in vivo showed the similarity as in vitro. The xenograft growth curve displayed that breast carcinoma with WISP/CCN5 gene knockdown possessed faster formation and development compared with negative group and control group (*P*<0.05) (Figure 3). However, there was no significant difference between null vector group and control group (*P*<0.05).

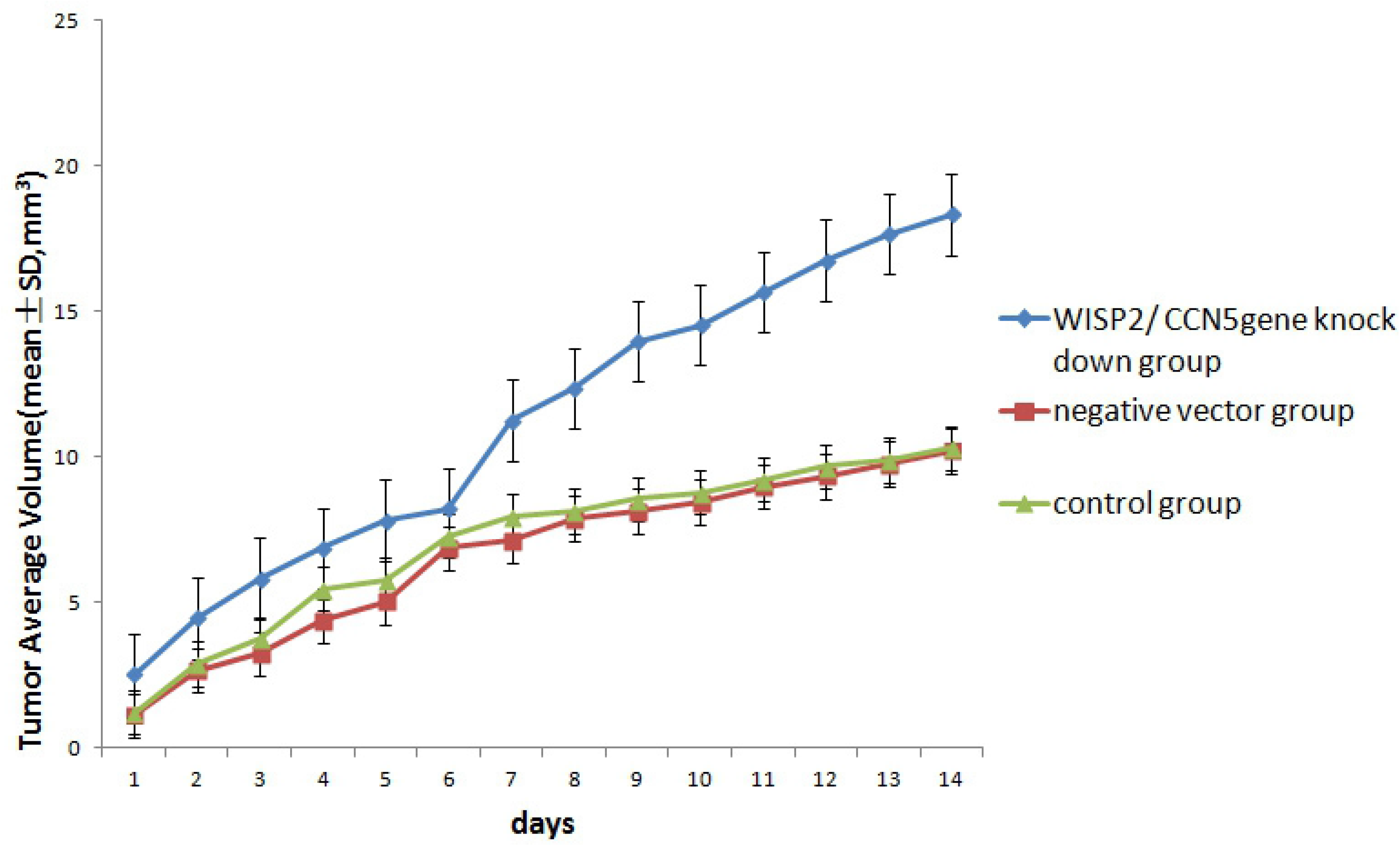
tumor growth curve.

### Knockdown of *WISP2/CCN5* gene resulted in abnormal level in mRNA and protein of Skp2 and p27Kip1 in vitro

RT-PCR analysis revealed that increased *Skp2* and decreased *p27Kip1* was observed in WISP2/CCN5shRNA group as compared to control group (*P*<0.05) as shown in Figure 4A. However, there was no marked difference between control group and negative control group (*P*>0.05). These experimental results suggested that up regulation of *Skp2* mRNA and down regulation of *p27Kip1* mRNA may be due to knockdown of *WISP2/CCN5*.

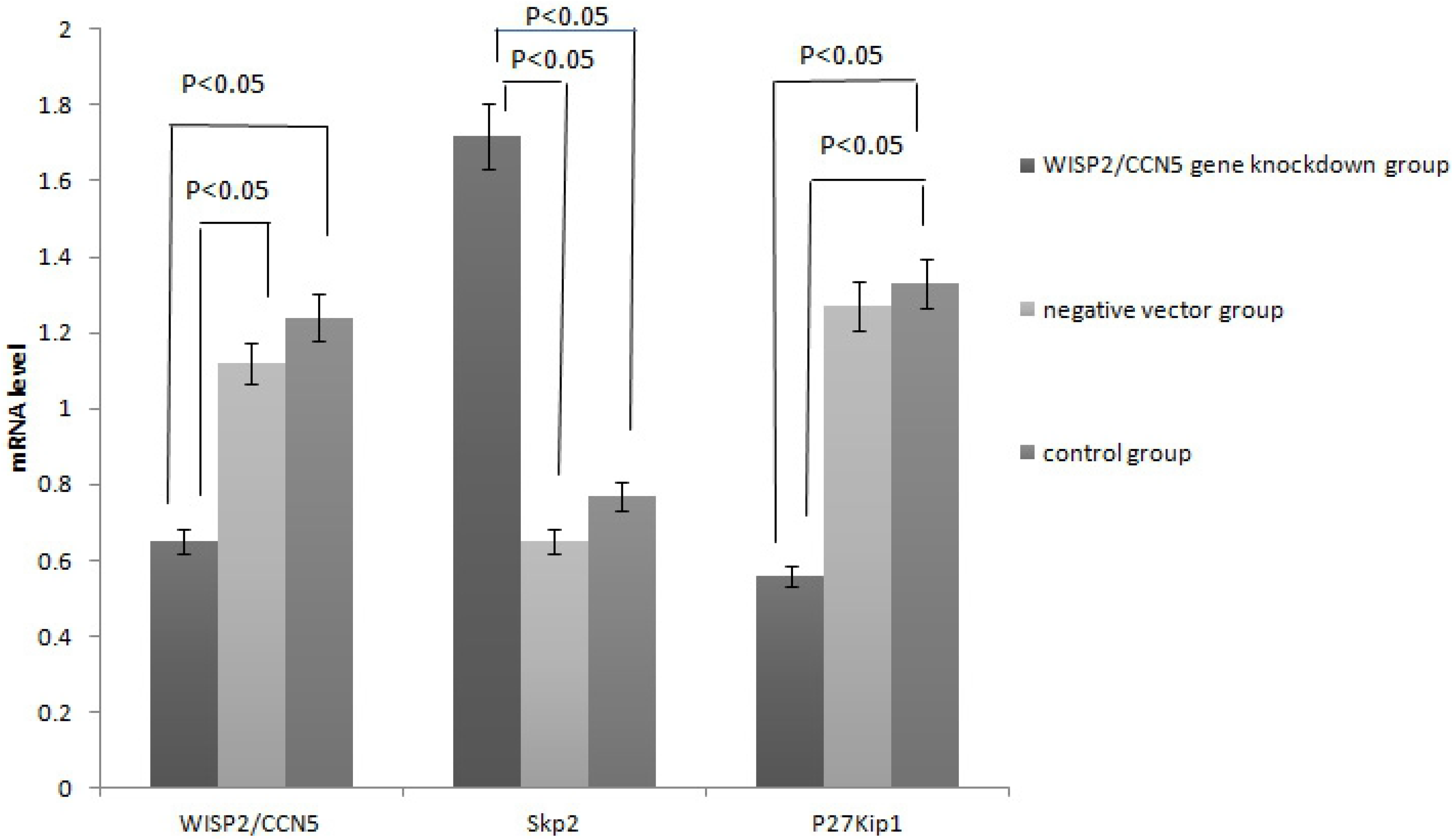
MCF-7 RT-PCR.

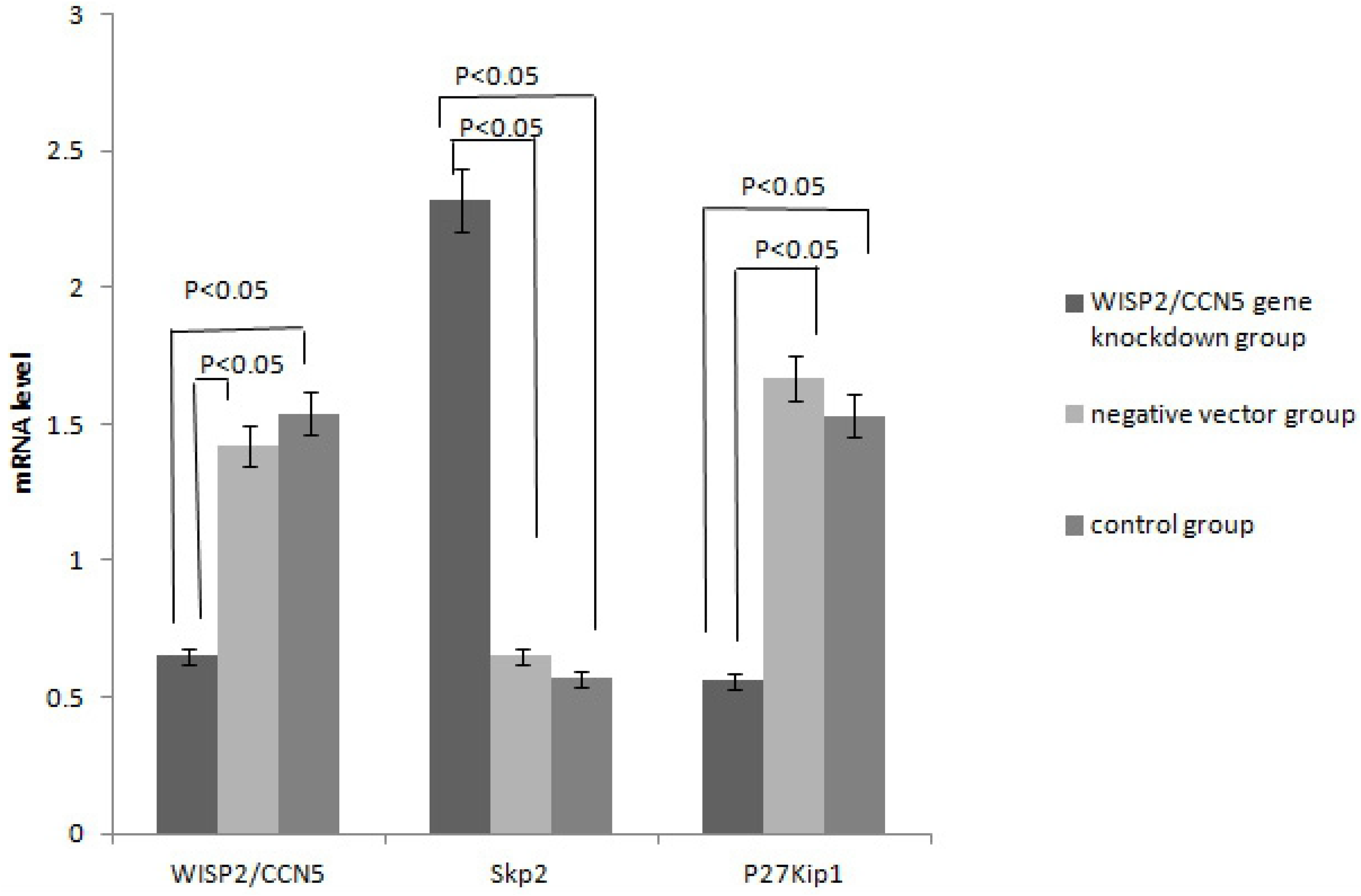
RT-PCR in xenografts.

In addition, western blot analysis presented rising Skp2 and declining p27Kip1 in WISP2/CCN5 gene knockdown group compared with negative vector group and control group(*P*<0.05)(Figure5B). Consequently, western blot stripe of Skp2 displayed darker and wider in WISP2/CCN5 gene knockdown group than negative vector group and control group, while lighter and thinner stripe of p27Kip1 was showed in WISP2/CCN5 gene knockdown (Figure 5A).

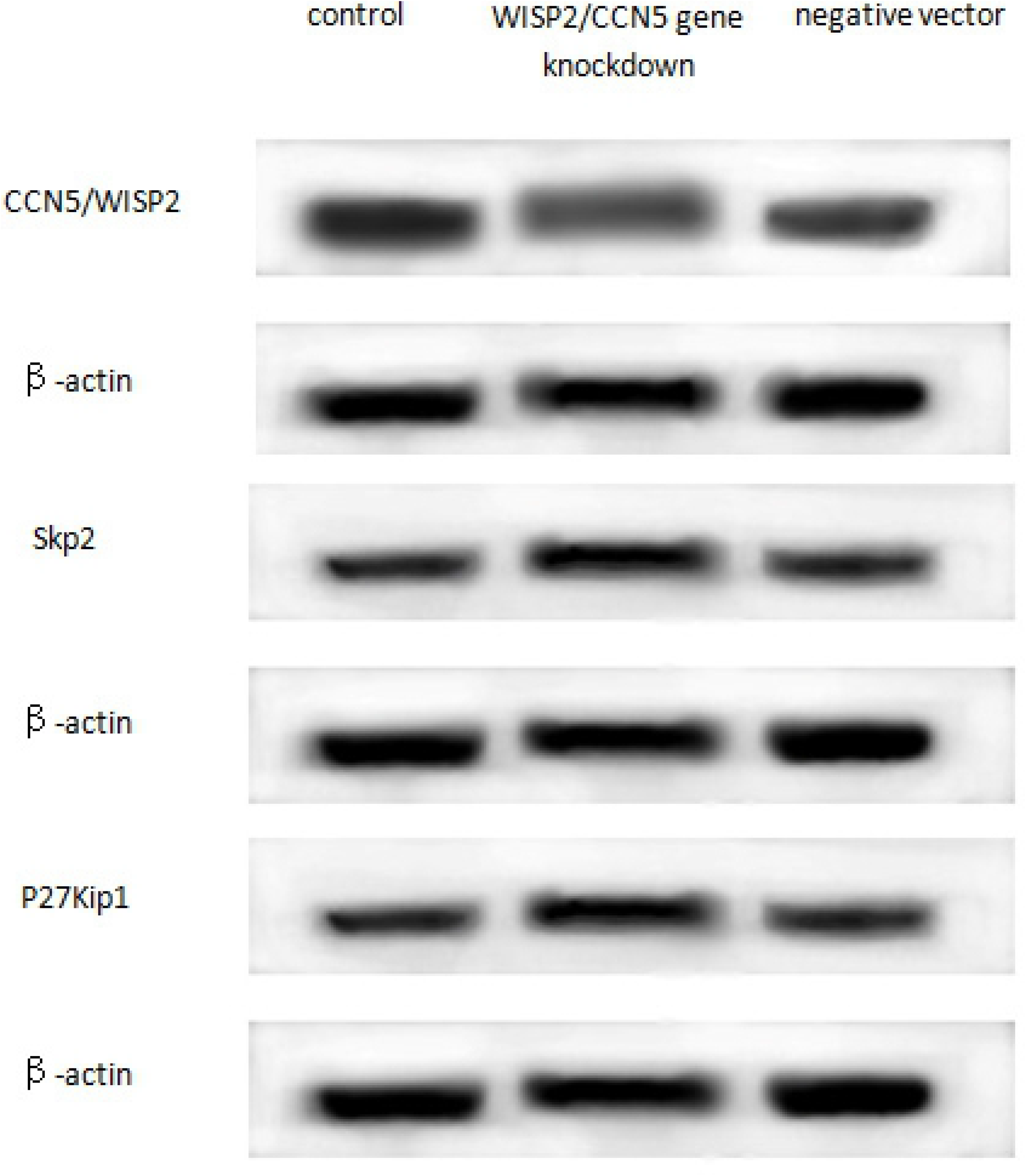
western blotting analysis.

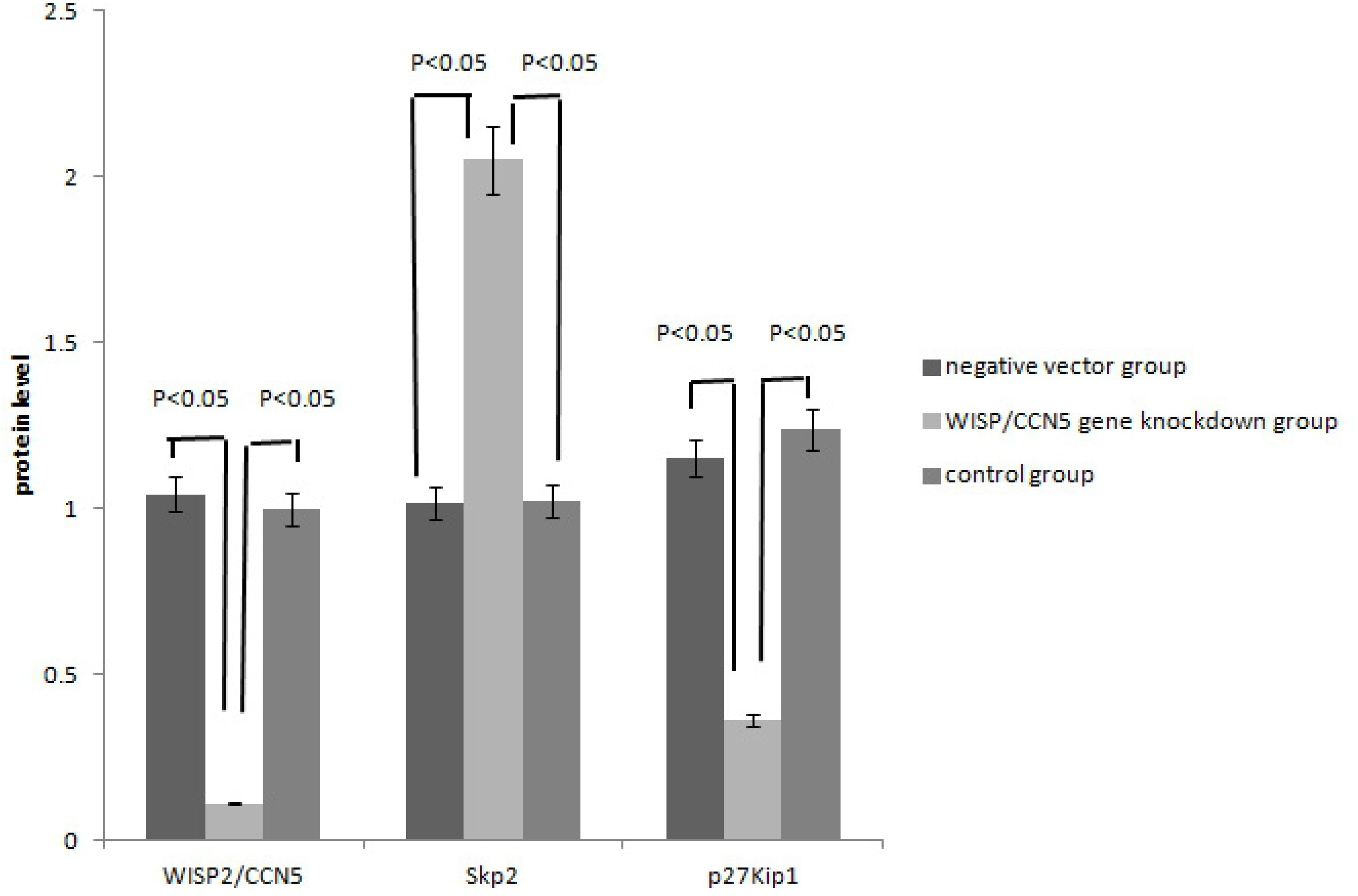
MCF-7 western blot.

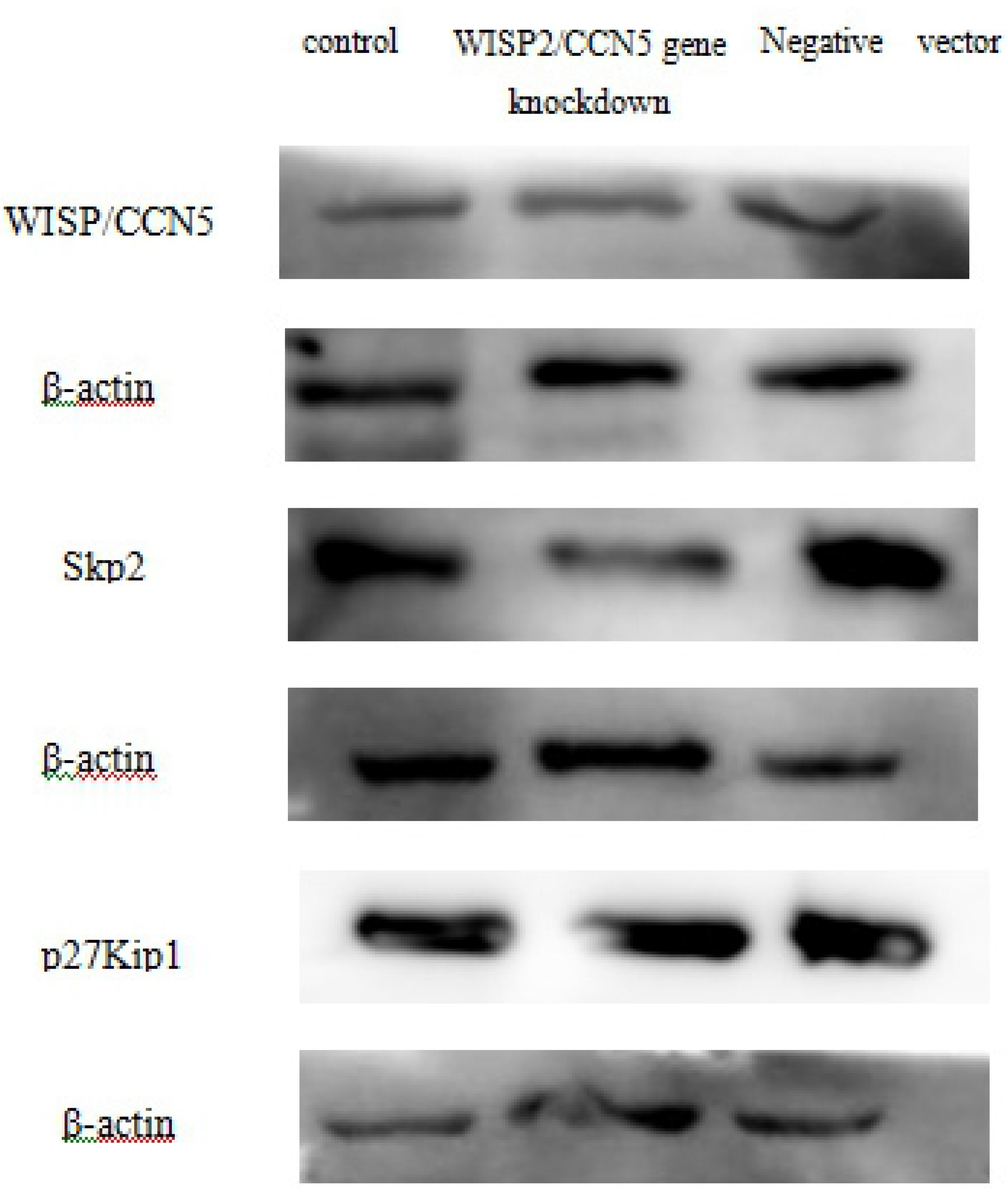
westernblotting.

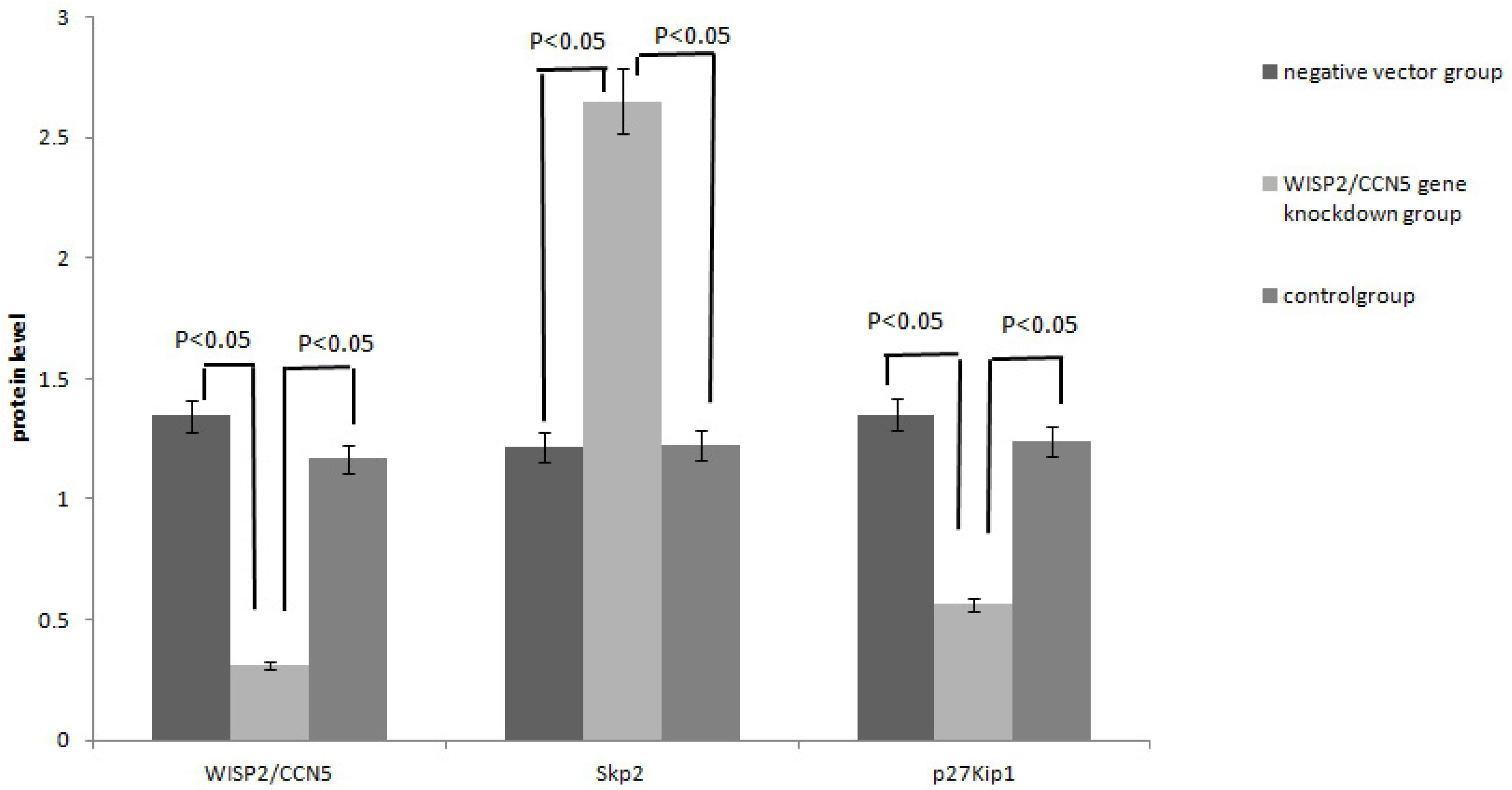
xenograft western blot.

The results of correlation analysis between WISP2/CCN5 and Skp2 and p27Kip1 were showed in Table 1 and Table 2. There was significant correlation between WISP2/CCN5 and Skp2 and p27Kip1 at mRNA and protein levels in three groups, including *WISP2/CCN5* gene knockdown group, negative vector group and control group. Especially, correlation in *WISP2/CCN5* gene knockdown group was remarkable between WISP2/CCN5 and Skp2 and p27Kip1 at mRNA and protein levels in *WISP2/CCN5* gene knockdown group.

**Table 1.**
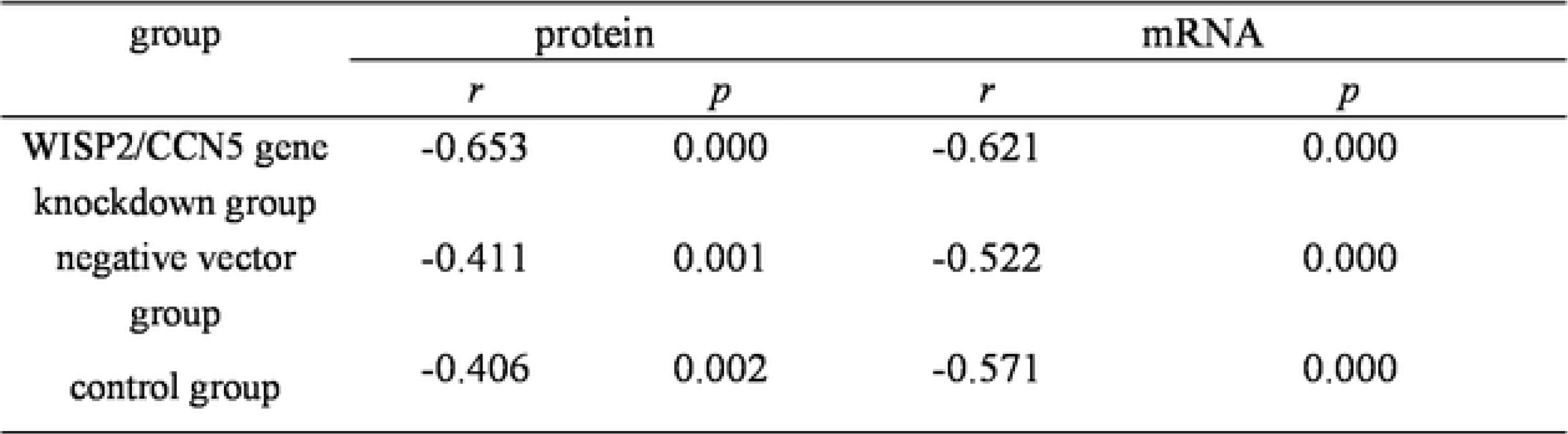
Correlation analysis between W1SP2/CCN5 and Skp2 in MCF-7 cells

**Table 2.** Correlation analysis between Skp2 and p27Kipl in MCF-7 cells

| group | protein |  | mRNA |  |
| --- | --- | --- | --- | --- |
|  | *r* | *p* | *r* | *p* |
| WISP2/CCN5 gene knockdown group | -0.653 | 0.000 | -0.621 | 0.000 |
| negative vector group | -0.411 | 0.001 | -0.522 | 0.000 |
| control group | -0.406 | 0.026 | -0.571 | 0.000 |

### Knockdown of *WISP2/CCN5* gene resulted in abnormal level in mRNA and protein of Skp2 and p27Kip1 in vivo

In the animal experiment, knockdown of *WISP2/CCN5* gene elevated the Skp2 and inhibited p27Kip1 at mRNA and protein level. The experiment results were showed in figure 4B and figure 5(C-D).

As shown in figure 4B of RT-PCR results analysis, increased *Skp2* mRNA level emerged in *WISP2/CCN5* gene knockdown group, while *p27Kip1* mRNA level declined (*P*<0.05), by contrast, in negative vector group and control group, there was no remarkable change (*P*>0.5).

At the same time, western blot analyses of Skp2 and p27 Kip1 in paired *β* -actin indicated up regulation of Skp2 protein level and down regulation of p27Kip1 protein level due to *WISP2/CCN5* gene knockdown(*P*<0.05) (Figure 5D). Similar changes were observed in western blot stripes (Figure 5C). Skp2 protein blot appeared darker stripe in *WISP2/CCN5* gene knockdown group than negative vector group and control group. However, p27Kip1 protein blot presented lighter stripe in *WISP2/CCN5* gene knockdown group.

The results of correlation analysis between WISP2/CCN5 and Skp2 and p27Kip1 were showed in Table 3 and Table 4. There was significant correlation between WISP2/CCN5 and Skp2 and p27Kip1 at mRNA and protein levels in three groups, including *WISP2/CCN5* gene knockdown group, negative vector group and control group. Especially, correlation in *WISP2/CCN5* gene knockdown group was remarkable between WISP2/CCN5 and Skp2 and p27Kip1 at mRNA and protein levels in *WISP2/CCN5* gene knockdown group.

**Table 3.** Correlation analysis between W1SP2/CCN5 and Skp2 in Xenografts

| group | protein |  | mRNA |  |
| --- | --- | --- | --- | --- |
|  | *r* | *p* | *r* | *p* |
| WISP2/CCN5 gene knockdown group | -0.732 | 0.000 | -0.765 | 0.000 |
| negative vector group | -0.452 | 0.001 | -0.574 | 0.000 |
| control group | -0.401 | 0.002 | -0.523 | 0.000 |

**Table 4.**
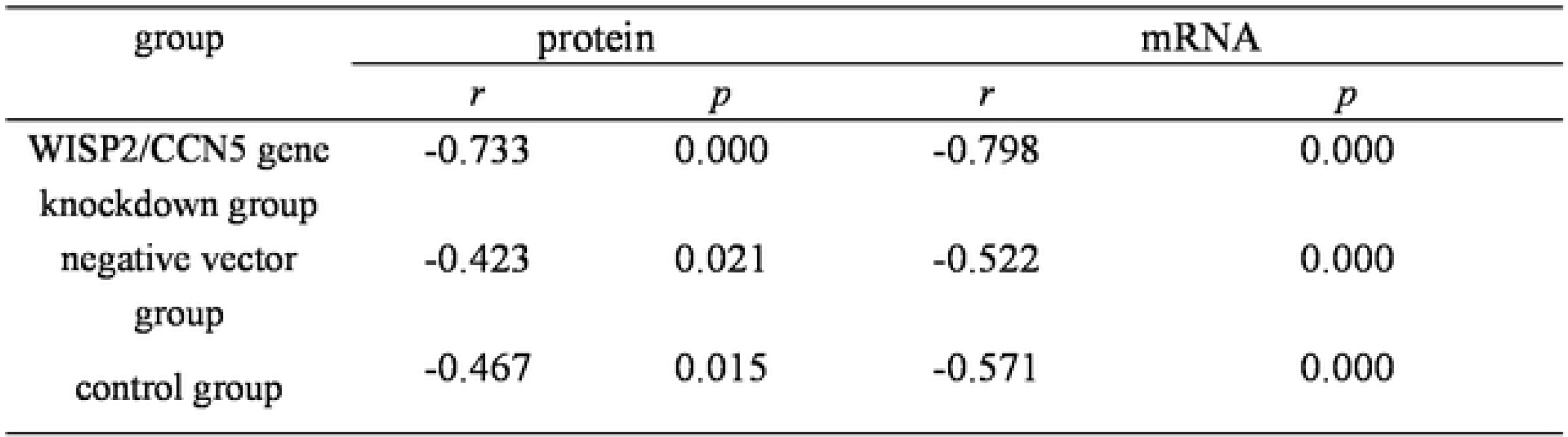
Correlation analysis between Skp2 and p27Kip 1 in Xenografts

## Discussion

Up to now, the role of WISP2 in ESCC is unelucidated, although the function of WISP2/CCN5 was explored in a variety of human cancers ^[8–10]^.

Our studies provide two significant discoveries. **First, our findings proved that deficiency of WISP2/CCN5 promote Skp2 expression via p27Kip1.** Skp2 and p27Kip1 in mRNA and protein levels were remarkably altered in *WISP2/CCN5* gene knockdown group. In particular, Skp2 mRNA expression was higher in *WISP2/CCN5* gene knockdown group than control group, whereas p27Kip1 mRNA expression was lower in *WISP2/CCN5* gene knockdown group than control group. Thus, these data demonstrated a significant relationship between *WISP2/CCN5, Skp2* and *p27Kip1* at the transcriptional level in breast cancer. These results demonstrated that aberrant level of *WISP2/CCN5* mRNA resulted in abnormal *Skp2* mRNA and subsequent unusual *p27Kip1*mRNA. Loss of *WISP2/CCN5* gene may promote breast cancer oncogenesis through *Skp2* and *p27Kip1*at mRNA level. A previous report showed that WISP2/CCN5 expressions in glioma cells possibly contributed to slower glioma cell proliferation ^[11]^. In addition, our current findings through western blotting showed Skp2 protein levels were increased, whereas p27Kip1 protein levels were decreased in *WISP2/CCN5* gene knockdown group. These results showed that loss of WISP2/CCN5 protein resulted in over expression of Skp2 protein and decreased level of p27Kip1 protein expression in MCF-7 cells and xenografts. Altogether, **these results suggested that WISP2/CCN5 resulted in down regulation of Skp2 and up regulation of p27Kip1.** Actually, the relationship of WISP-2/CCN5 expression with less frequent invasive phenotype of breast carcinoma has been explored by some researchers ^[12–13]^, S-phase kinase-associated protein 2 (SKP2) is an oncogene and cell cycle regulator that specifically recognizes phosphorylated cell cycle regulator proteins and mediates their ubiquitination, which is overexpressed in lymphoma^[14]^, prostate cancer^[15]^, melanoma^[16]^, nasopharyngeal carcinoma ^[17]^, breast cancer^[18]^.

P27Kip1, as one of the best known protein substrates of Skp2, is the cyclin-dependent kinase (CDK) inhibitor 1B (CDKN1B)^[15]^. Moreover, over expression of Skp2 and low expression of p27Kip1 are strongly associated with aggressive tumor behavior and poor clinical outcomes in a variety of cancers ^[16–18]^.

**Second, our studies provided the new evidence showing WISP2/CCN5 as a tumor suppressor in breast cancer in vitro animal assay and in vivo breast cancer cells through Skp2 via p27Kip1.** In the present study, we reported that *WISP2/CCN5* gene knockdown induced proliferation of MCF-7 cell line and growth of breast carcinoma, and aberrant mRNA and protein level of Skp2 and p27Kip1. Mechanistically, WISP2 exerts its tumor suppressive functions via regulation of Skp2 and p27Kip1 in breast cancer.

WISP2/CCN5 has been reported to govern several gene expressions in human cancer cells. For instance, deficiency of WISP2/CCN5 promotes mesenchymal to epithelial transition (MET) ^[21]^. Moreover, MET is required for the growth of micrometastatic breast cancer ^[22]^. **Our study reported that** *WISP2/CCN5* **gene knockdown may promote the proliferation of breast cancer cells and development of breast carcinoma.** It was ever reported that loss of WISP2/CCN5 in MCF-7 breast cancer cells can also promote the emergence of a cancer stem-like cell phenotype characterized by high expression of CD44^[23]^. That demonstrated that loss of WISP2/CCN5 play an important role in tumorigenesis, in another world WISP2/CCN5 inhibit the carcinogenensis. Simultaneously, some researchers also hold the same opinion as us that loss of WISP2/CCN5 protein in MCF-7 promotes a stem-like cell, that means deficiency of WISP2/CCN5 have closed relationship with monoclonal proliferation of breast cancer cells ^[24–26]^. Consequently, the role of WISP2/CCN5 in development of breast cancer has received widespread attention ^[27–29]^.

**Finally**, **we believed that underlying mechanisms of MCF-7 cells proliferation and tumorigenesis may be results of decreased WISP2/CCN5 through aberrant levels of Skp2 and p27Kip1.** There was ever research reported that induced activation of CCN5 in triple-negative breast cancer (TNBC) cells promotes cell growth arrest at the G0/G1 phase, reduces cell proliferation and delays tumor growth in the xenograft model^[30]^, whereas our research showed that the regulation between WISP-2/CCN5 and Skp2 with p27Kip1 in MCF-7 cell line with ER positive.

**We highlighted that WISP2/CCN5 may be a promising therapeutic target and development of WISP2/CCN5 promoter would have a great impact on breast cancer therapy.** The relationship between deficiency of WISP2/CCN5 and estrogen dependent MCF-7 has been explored because WISP2/CCN5 is an estrogen response gene in ER-α-positive breast cancer cells, such as MCF-7 cell line, and present studies showed that the estradiol-treatment is able to activate CCN5 transcription ^[31–33]^. Therefore, correlation between WISP2/CCN5 and ER, PR and Her-2 still need to be detected in breast cancer.

## Acknowledgments

We thanks for the support from Science Program of Education Committee, Tianjin,China (No.2017 KJ160).

